# Highly enhanced transgene integration in human pluripotent stem cells by serine integrase BxbI

**DOI:** 10.1101/2024.10.18.619063

**Authors:** Matias I. Autio, Arnaud Perrin

**Author notes:** corresponding author &.

## Abstract

Genome engineering and especially integration of large DNA payloads in human pluripotent stem cells (hPSC) remains challenging largely due to the sensitive nature of hPSCs. Site specific recombinases such as the serine integrase Bxb1 have been utilised successfully to target hPSC, however the targeting efficiencies remain low. Here we leveraged codon optimisation, inclusion of a nuclear localisation signal, mRNA modality and co-introduction of p53 dominant negative fragment to increase the transgene payload integration efficiency by up to 20-fold in defined genomic safe harbours in hPSCs.

## Introduction

Modern genome engineering technologies have enabled modification of mammalian cells in a targeted manner from single nucleotide changes to insertions of larger transgenic payloads. However, methods utilising targeted nucleases, such as CRISPR/Cas9, rely on the cellular DNA repair machinery for homology directed repair mediated integration, require the generation of exposed DNA double-stranded breaks (DSB), suffer from very low efficiency when inserting large DNA cargoes and frequently result in unwanted editing outcomes^1–4^. Human pluripotent stem cells (hPSCs) offer great potential for cell therapy applications through integration of therapeutic payload genes and differentiation to the required cell type^5–8^ however hPSCs are particularly difficult to engineer due to their susceptibility to apoptosis induced by p53-mediated DNA damage response^9,10^ . Serine integrases, such as BxbI, do not rely on the cellular machinery and are not thought to generate exposed DSB, thus they have recently been used to successfully integrate large payloads into hPSCs, both directly into pre-engineered landing pads ^11–13^ and in conjunction with Cas9-mediated targeting^14–16^. It is possible to generate 100% engineered hPSC populations through the integration of selection markers^11,17,18^, however without selection the reported targeting efficiencies remain very low often below 1% ^13,18^. Many applications would benefit from higher integration efficiency of large payloads in hPSCs, thus there have been efforts to evolve a more efficient BxbI protein, however these efforts have so far given rise to only 3.8% efficiency in hPSC targeting^16,19^. In this study, we set out to test whether we can increase the targeting efficiency of the BxbI integrase in hPSCs through optimisation of the nucleotide sequence, method of delivery and inhibition of the p53-pathway.

## Methods

### Plasmid construction

All restriction enzymes were purchased from NEB. PCR reactions were conducted using Q5® Hot Start High-Fidelity 2X Master Mix (NEB, M0494L). Ligations were conducted using isothermal assembly with NEBuilder® HiFi DNA Assembly Master Mix (NEB, E2621L). All gBlocks were ordered from Integrated DNA Technologies, Singapore.

Sequences encoding the BxbI integrase linked with or without C- and N-terminal bi-partite nuclear localisation signal (BP-NLS) were codon optimised using iCodon online tool^20^. The optimised sequences were used to design gBlocks with appropriate overhangs (Fig. 1-3) and cloned into pMAX-GFP plasmid from Lonza, digested with KpnI and SacI (Fig. 4, Supp. File 1).

**Figure 1.**
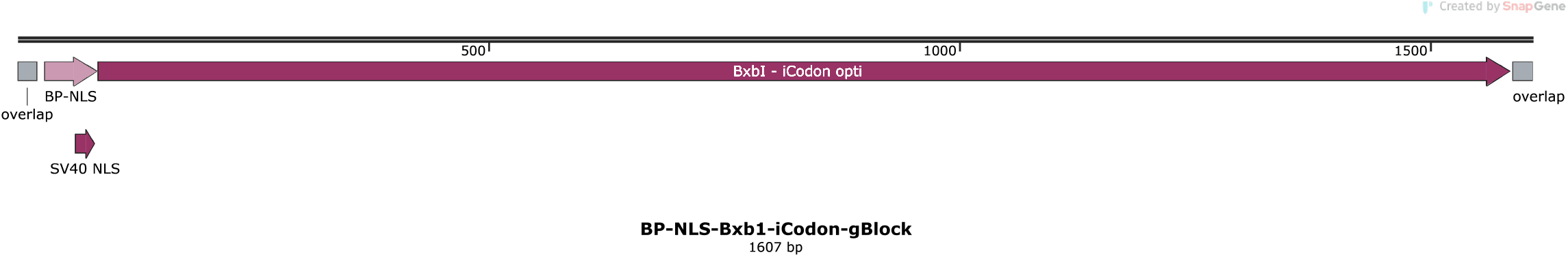
Schematic representation of BP-NLS-BxbI-iCodon gBlock

**Figure 2.**
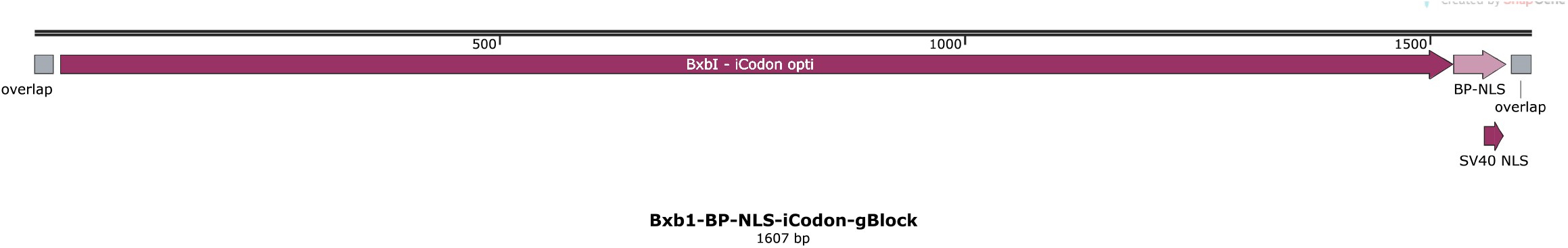
Schematic representation of BxbI-BP-NLS-iCodon gBlock

**Figure 3.**
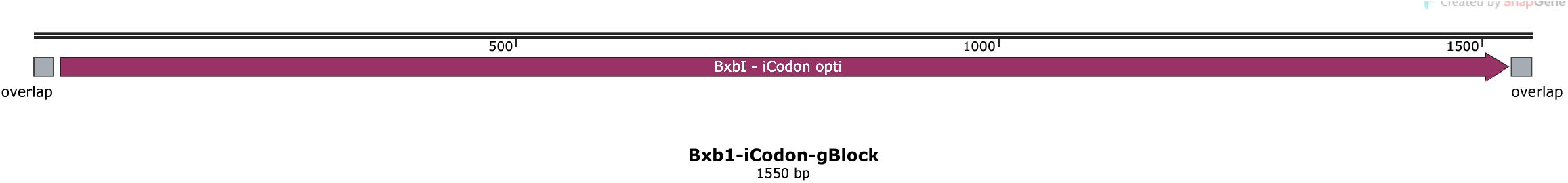
Schematic representation of BxbI-iCodon gBlock

**Figure 4.**
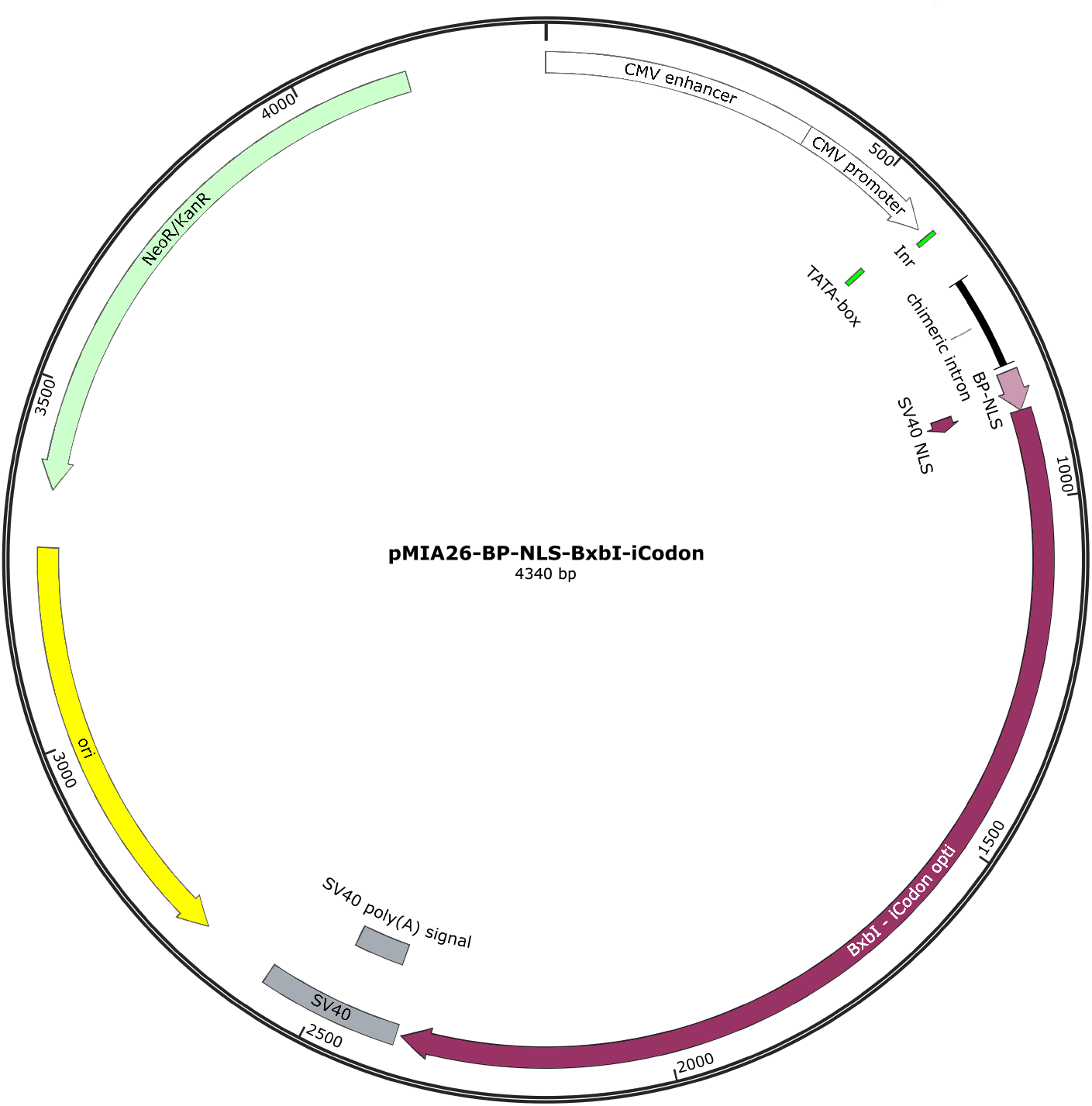
Schematic representation of pMIA26 plasmid

Donor payload plasmid pMIA10.53_copGFP_iC-opti (Supp. File 2) was ordered from Twist Biosciences (San Fransisco, CA, USA).

### mRNA sequence design

The mRNA sequences corresponding to the coding sequence of pMIA22^13^, the BP-NLS-BxbI (pMIA26) optimised for this study, tp53 dominant negative fragment (tp53dn) sequence from pMIA3^21^ as well as p53dn-P2A-BP-NLS-BxbI sequence optimised with iCodon, were all flanked by Xenopus globin 5’-UTR and 3’-UTR and synthesised by Azenta/GENEWIZ (Burlington, MA, USA) with pseudouridine and Cap1 (Fig. 5-8, Supp. File 3-6)

**Figure 5.**
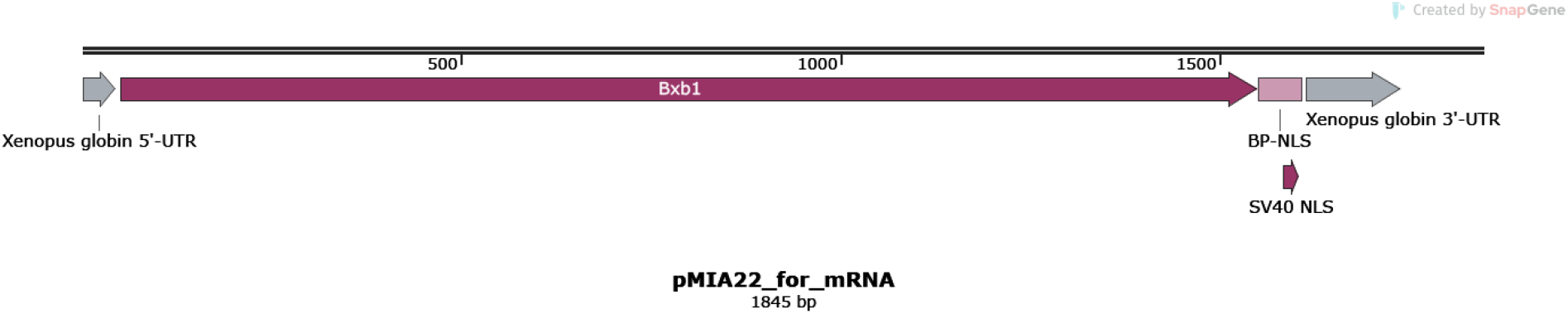
Schematic representation of pMIA22 mRNA

**Figure 6.**
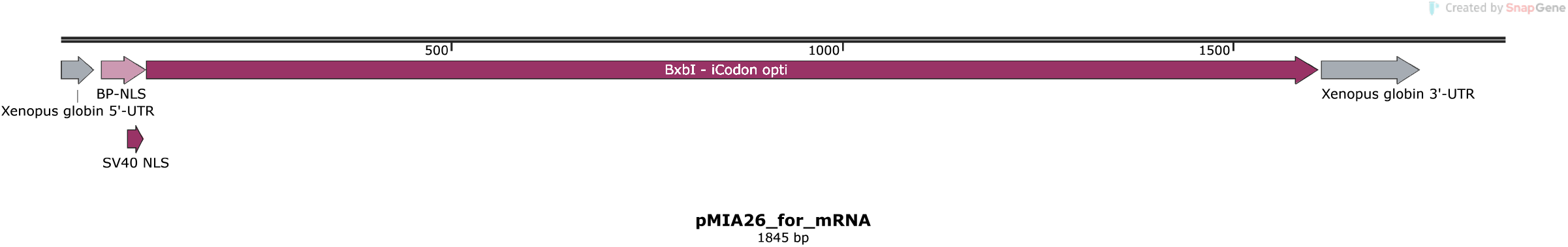
Schematic representation of pMIA26 mRNA

**Figure 7.**
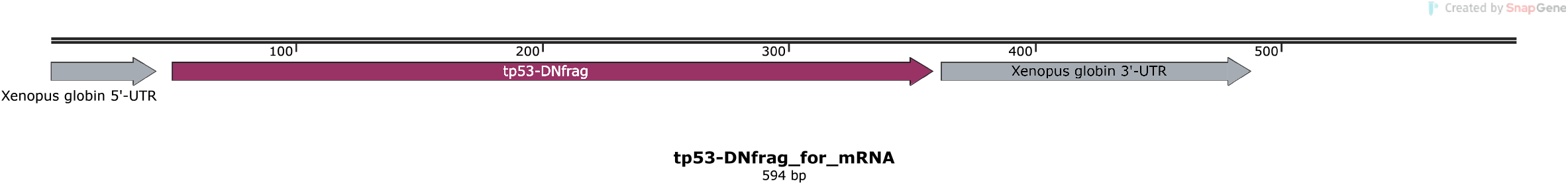
Schematic representation of tp53dn-fragment mRNA

**Figure 8.**
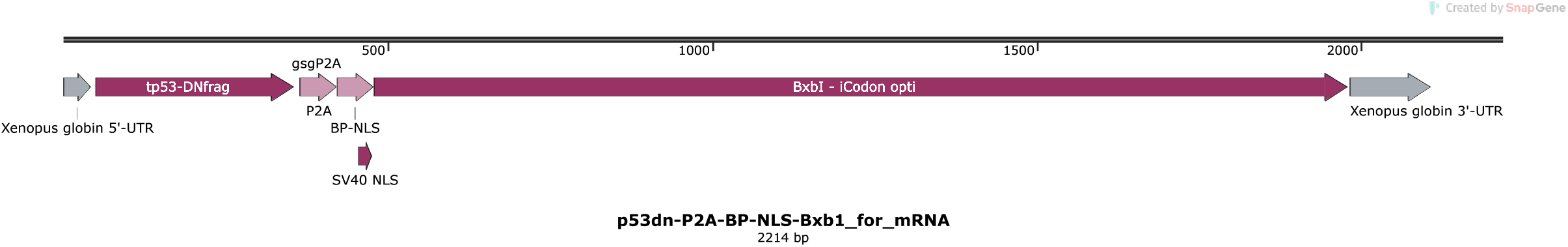
Schematic representation of p53dn-P2A-pMIA26 mRNA

### Stem cell culture

Human ESC lines H1 (WiCell, WA01) & H9 (WA09) engineered to contain a BxbI specific landing-pad expression cassette at the Pansio-1 safe harbour locus^13^ (Supp. File 7) were maintained using mTeSR medium (STEMCELL Technologies, 85850) on 1:200 Geltrex (ThermoFisher Scientific, A1413202) coated tissue culture plates and passaged regularly as cell aggregates every 4-5 days using RelesR (STEMCELL Technologies, 05872). Identity of H1 cells was authenticated by the supplier, WiCell. Identity of H9 cells was authenticated by short tandem repeat analysis (AxilScientific). The cell cultures were tested for mycoplasma contamination monthly and confirmed negative.

### hPSC targeting with Amaxa Nucleofector

Five μg of plasmids encoding different versions of BxbI (pMIA22, pMIA26 and pCMV-BxbI, Addgene #51552^22^ a kind gift from Dr. Michele Calos) each together with five μg of the pMIA10.53 donor plasmids were nucleofected into hESC using the P3 Primary Cell kit (V4XP-3024) and programme CA-137. 1.5×10^6 cells were used for each targeting and were plated onto geltrex coated wells on 6-well plates in mTeSR with CloneR2 (STEMCELL Technologies, 100-0691) following nucleofection. After 24h media was changed to mTeSR, and cells were allowed to recover for another 24-48h before analysis.

### hPSC targeting with Lipofectamine Stem reagent

For each targeting reaction 2.5 μg of pMIA10.53 donor plasmid was mixed into 125 μl of Optimem (ThermoFisher, 31985070) along with either 2.5 μg DNA plasmid recombinase, 2.5 μg mRNA recombinase or 1.25 μg of mRNA recombinase and 1.25 μg of mRNA tp53-dn. 20 μl of Lipofectamine Stem (ThermoFisher, STEM00015) reagent was mixed in 125 μl of Optimem and combined with the Optimem aliquot containing the targeting constructs. The transfection mixture was left to incubate at room temperature for 10 min. At the same time hPSCs were prepared into single cell suspension using Accutase (ThermoFisher, A1110501). After neutralisation and counting the cells were aliquoted into 500 000 cells per targeting. The transfection mixture was used directly to resuspend the spun-down aliquot of 500 000 cells. After resuspension the cells and the mixture were plated onto geltrex coated wells containing mTeSR supplemented with CloneR2 and left to recover for 24h. After 24h media was changed to mTeSR, and cells were allowed to recover for another 24-48h before analysis.

### Flow cytometry analysis

H1 & H9 Pansio-1 lines targeted with different BxbI expressing modalities and pMIA10.53-copGFP donor were disassociated with accutase (STEMCELL Technologies, 07922) and resuspended in PBS. The single cells in PBS were analysed at Flow Cytometry Core, SIgN A^*^STAR using a BD FACSymphony A3 Analyzer and FlowJo (v10.8.1).

## Results

Integration of large DNA payloads into hPSCs remains a challenging task. In our hands, efficiency of targeting with previously published BxbI and donor constructs^13,22^ into established hPSC landing pad lines^13^ remains at maximum around 2.5% (Fig. 9). To investigate the effect of nuclear localisation signal positioning and codon optimisation on BxbI payload integration we designed three new versions of BxbI using the iCodon online tool^20^: BxbI-iC, BP-NLS-BxbI-iC and BxbI-BP-NLS-iC (Fig. 1-4). We tested the new BxbI constructs alongside previously published pCMV-BxbI and pMIA22 constructs by nucleofecting the integrase constructs with a pMIA10.53-copGFP construct into H1-Pansio-1 cells. We observed successful targeting with four of the tested constructs. Interestingly the BxbI-iC construct did not show any increase in GFP positive cells over untargeted cells (Fig. 9). Targeting with BxbI-BP-NLS-iC and the previously published constructs, pCMV-BxbI and pMIA22, all resulted in approximately 2-2.5% GFP positive cells, however targeting with the BP-NLS-BxbI-iC construct gave rise to approximately 5% GFP positive cells (Fig. 9). These results suggest that a N-terminal bi-partite NLS increases BxbI mediated payload integration up to 2-fold over the wild type BxbI protein, we named the BP-NLS-BxbI-iC plasmid pMIA26 (Fig. 4, Supp. File 1).

**Figure 9.**
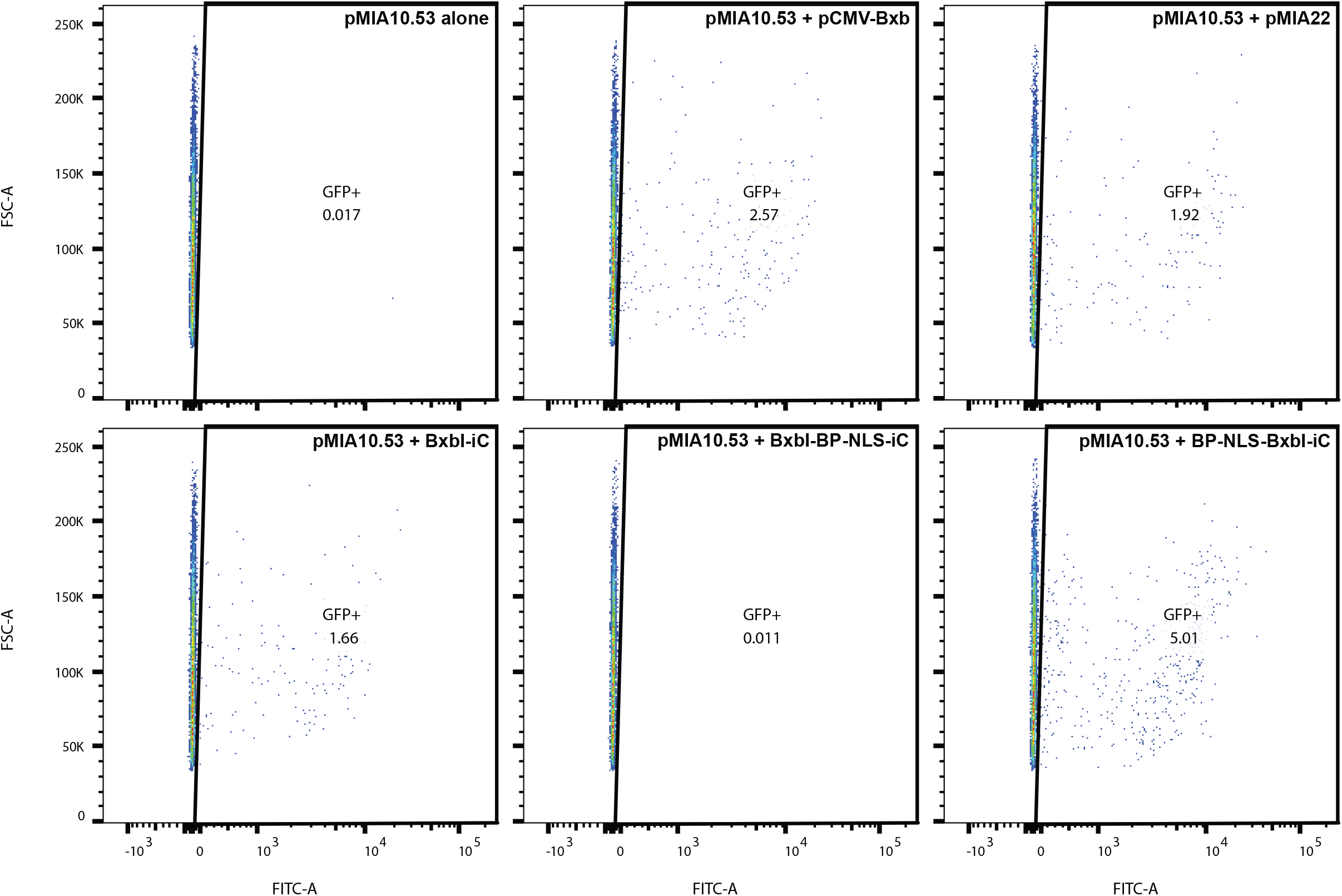
Scatter plots of flow cytometry analysis H1-Pansio-1 cells targeted with pMIA10.53 donor plasmid in combination with different BxBI variant plasmids. Gating on FSC-A and FITC-A is set with cells targeted with pMIA10.53 alone. Percentage of GFP positive cells is shown for each tested cell population.

Human PSCs are well known require specific conditions and careful handling to be maintained in culture^23^, furthermore hPSCs have a highly active p53-mediated DNA damage response^24^, both of which are explanations to significant numbers of cell death observed in treatment and targeting of hPSCs. Experiments investigating Cas9 mediated targeting of hPSCs further demonstrate the sensitivity of the cells to insults^9,10^. In our hands we frequently observed a substantial number of GFP positive cells detaching from the culture plates and dying following targeting with BxbI constructs (data not shown). These observations led us to explore ways to reduce the stress inflicted on the cells during the targeting. We designed mRNA versions of pMIA22 and pMIA26 BxbI constructs and in addition to these we designed a codon optimised mRNA sequence of tp53 dominant negative fragment^21^ individually or linked to the pMIA26 mRNA sequence (Fig.5-8, Supp. File 2-5) and targeted these to H9- and H1-Pansio-1 cells, alongside DNA versions of BxbI, using Lipofectamine Stem mediated reverse transfection. Using mRNA constructs of pMIA22 and pMIA26 we saw average increases of approximately 2-5-fold in the number of GFP positive cells compared to the DNA plasmid versions of the respective constructs (Fig. 10, Supp. File 7). Interestingly, targeting with mRNA of pMIA26 together with mRNA on tp53dn, in either separate or the same RNA sequence, led to further increase GFP positive cells over the mRNA of pMIA26 alone. The highest percentage of GFP positive cells was seen with pMIA26 and tp53dn mRNA expressed from separate sequences resulting in on average 21% positive cells, an increase of approximately 5-7-fold mRNA of pMIA26 alone (Fig. 10, Supp. File 8). Overall, the amount of hPSCs that integrated the GFP payload increased from ≤1%, using published DNA constructs, to ≥20% when using both mRNA and adding in the tp53dn fragment.

**Figure 10.**
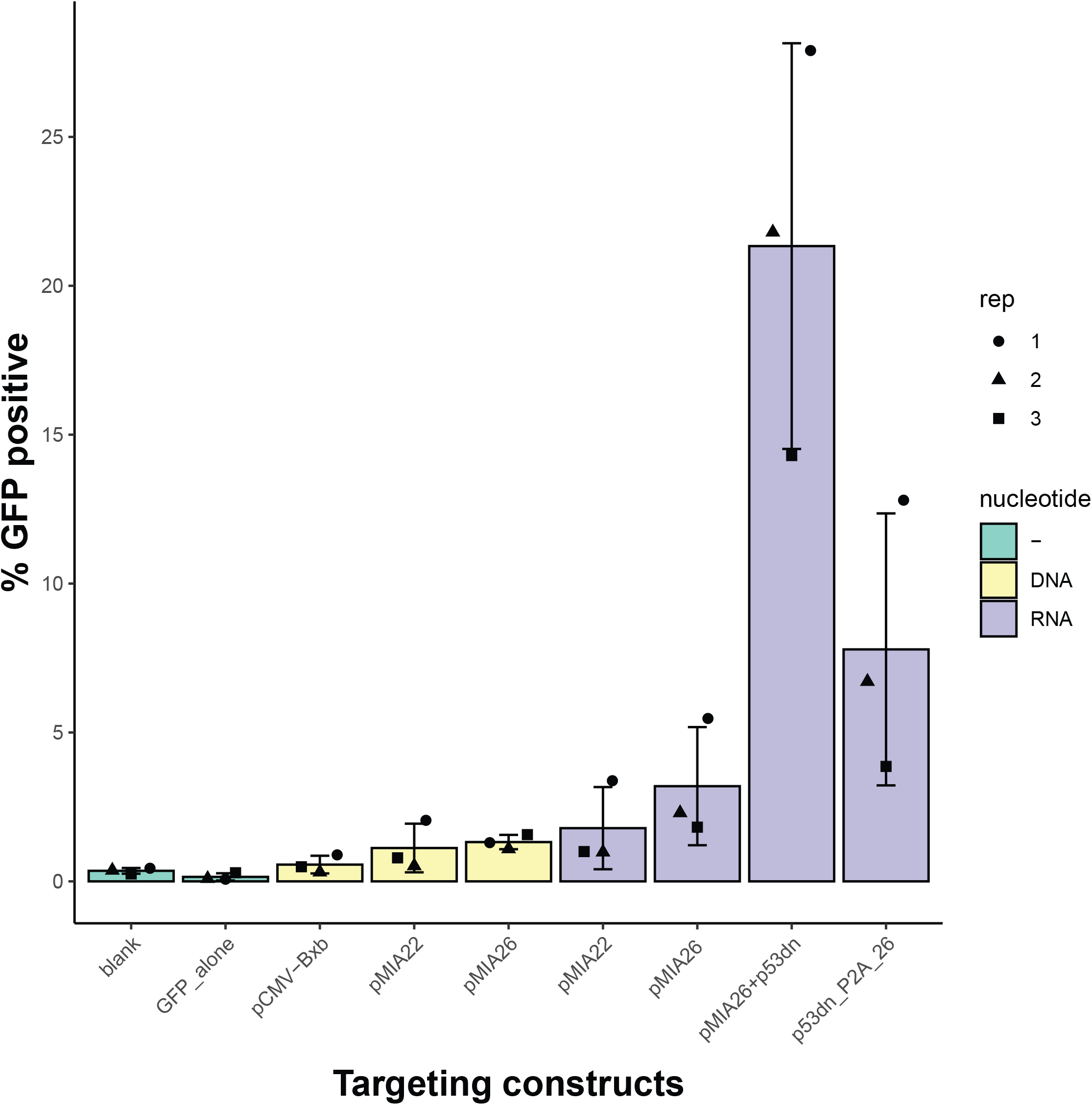
Fow cytometry analysis H9-Pansio-1 cells targeted with pMIA10.53-copGFP donor plasmid in combination with different BxBI variant constructs. Bar chart of average percentage of GFP positive cells over three replicate experiments is shown for each tested cell population. Error bars indicate standard deviation. Individual data points from each replicate experiment are shown.

## Discussion

Integration of large DNA payloads into hPSCs is of great interest for potential clinical applications with engineered therapeutic cells. However, efficiency of integration remains low, with even the most promising technologies such as serine integrases only reaching integration efficiencies in the low single digits^11,16,18^. There have been several recent publications that have used directed evolution to increase BxbI, the most well characterised serine integrase, integration efficiency mediated^16,19,25^. Only the eePASSIEGE system, with an evolved BxbI variant, has been tested in hPSCs and achieved approximately 3.8% payload integration efficiency. Here we demonstrate payload integration at ≥20% efficiency using a codon optimised wild type BxbI serine integrase linked with an N-terminal bi-partite NLS delivered in parallel with tp53dn-fragment both in mRNA format. Our results show a 5-fold increase over the highest published integration efficiency and up to 20-fold increase over published BxbI wildtype constructs transfected in a DNA format. The methodology for BxbI mediated payload integration reported here will enable high-throughput experiments, with e.g. targeted integration of single gRNAs for CRISPR-screens or targeted massively parallel reporter assays in hPSCs. The high efficiency payload integration will also address a bottle neck in genome engineering of hPSCs towards cell therapy. Combining the mRNA modality, BP-NLS and tp53dn-fragment with some of the recently reported evolved BxbI variants^16,19,25^ may enable even further gains in payload integration efficiency and will be of great interest to investigate further.

## Acknowledgements

We would like to express our gratitude to Dr Shyam Prabhakar and Dr Jay Shin for their generous support. The authors thank the Agency for Science, Technology and Research (A^*^STAR)’s Singapore Immunology Network (SIgN) Flow Cytometry platform for enabling the flow cytometry work for this publication. A^*^STAR’s SIgN Flow Cytometry platform is supported by SIgN, A^*^STAR, and the National Research Foundation (NRF), Immunomonitoring Service Platform (Ref: ISP: NRF2017_SISFP09) grant. This research was funded by A^*^STAR I&E GAP funding (I22D1AG046) to M.I.A.. The funding bodies played no role in the design of the study or in collection, analysis, and interpretation of data or in writing of the manuscript.

## Declaration of interests

M.I.A. and A.P. are listed as inventors on a patent application submitted on technology described in this manuscript (Singapore Patent Application 10202403256Y).

