## Supplementary figures and images for "Highly enhanced transgene integration in human pluripotent stem cells by serine integrase BxbI"

### Supplemental File 2

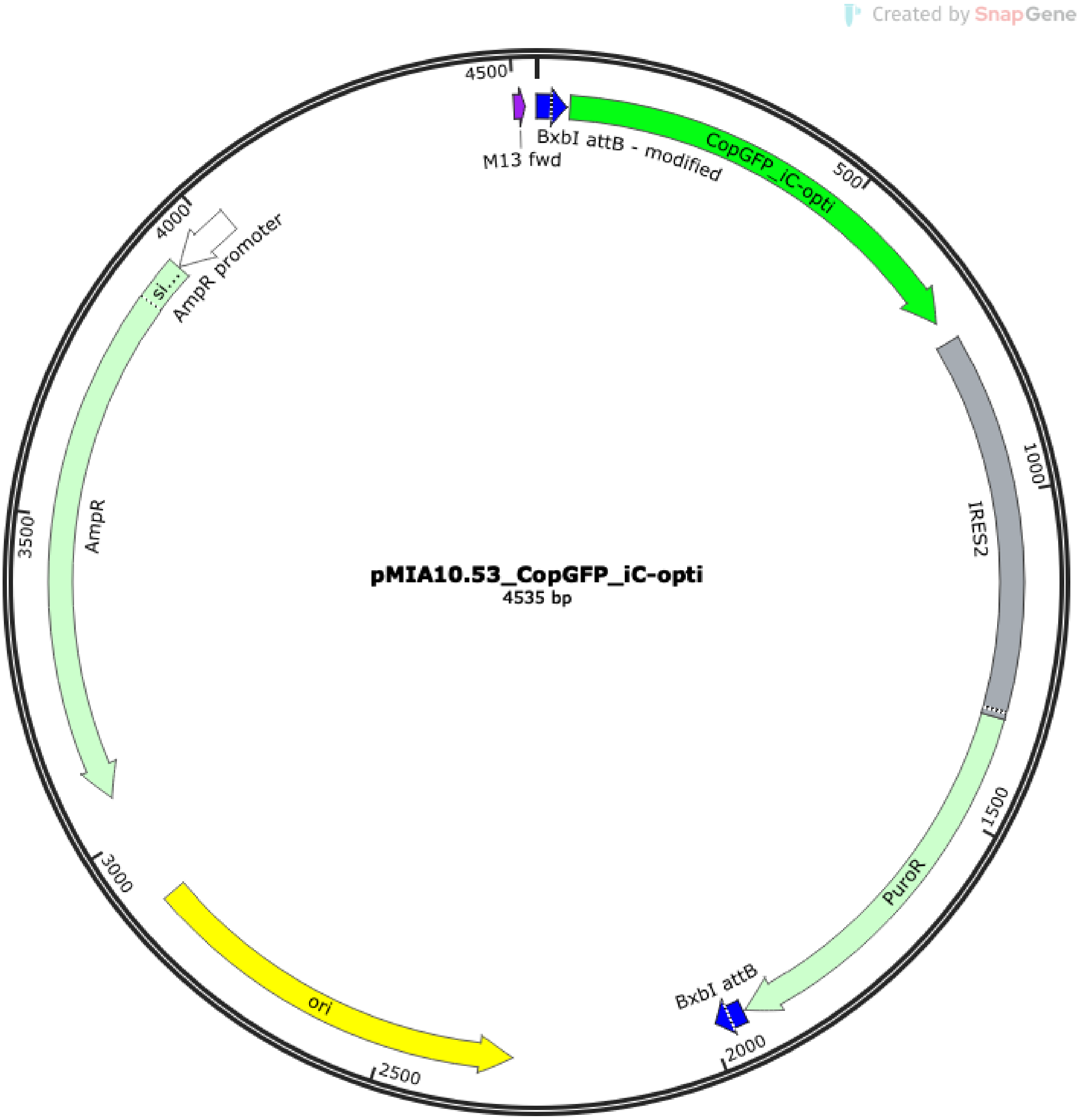


**Supplemental File 2**. Schematic representation of pMIA10.53_copGFP plasmid.

### Supplemental File 8

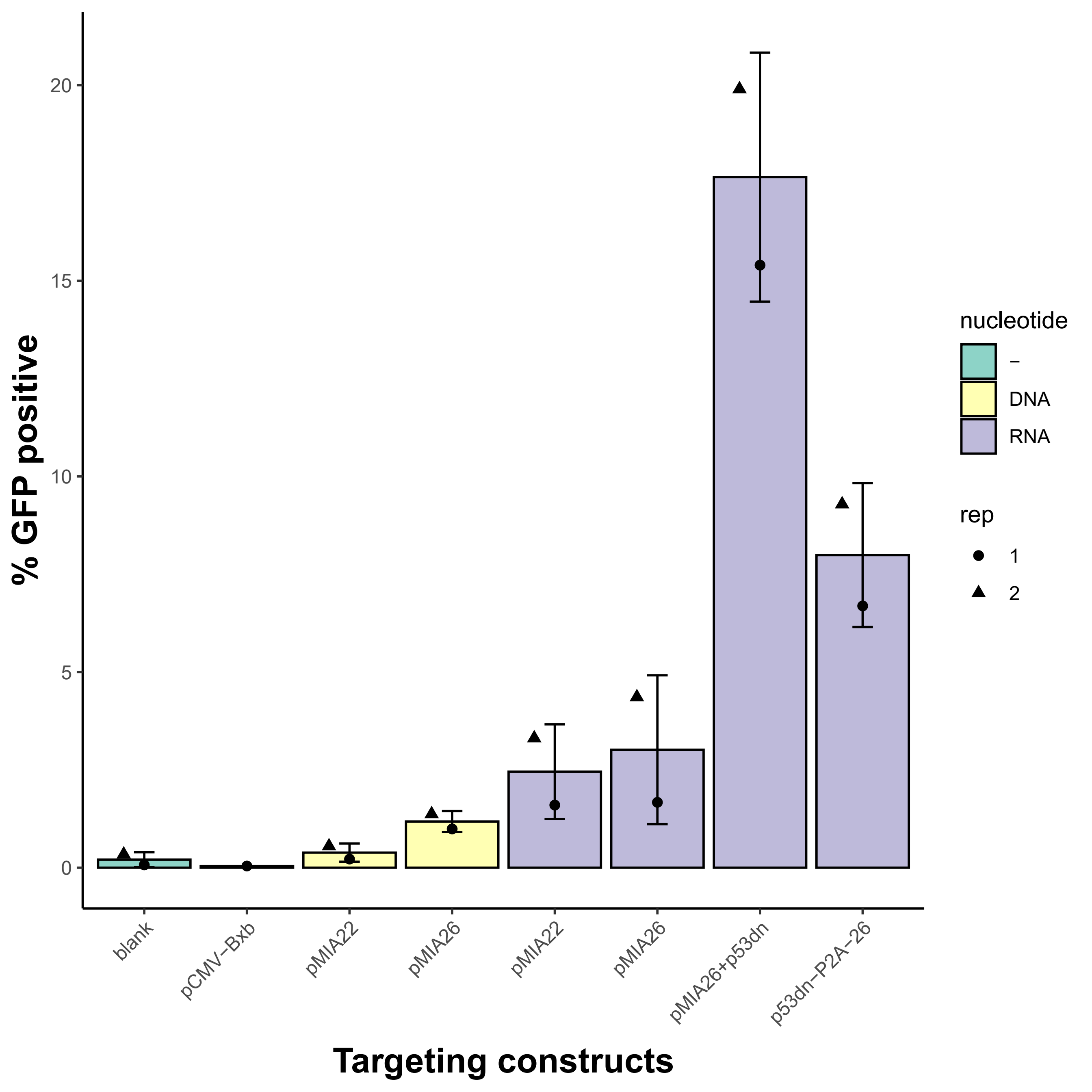
