## Supplemental File 7 for "Highly enhanced transgene integration in human pluripotent stem cells by serine integrase BxbI"

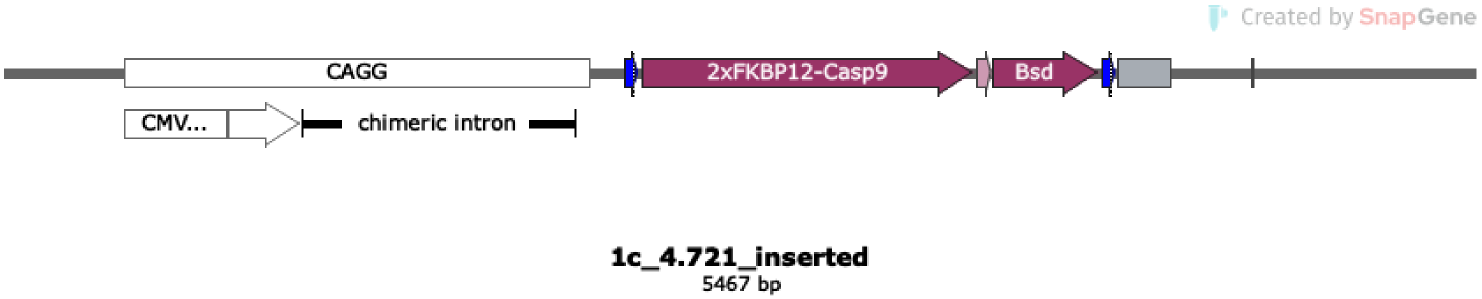


**Supplemental File 7**. Schematic representation of integrated landing pad expression cassette in H1-Pansio-1 cells. CAGG promoter in white. BxbI attP sites in blue. Protein coding sequences in maroon. P2A sequence in pink. PolyA sequence in grey
